# Stable but not rigid: Long-term *in vivo* STED nanoscopy uncovers extensive remodeling of stable spines and indicates multiple drivers of structural plasticity

**DOI:** 10.1101/2020.09.21.306902

**Authors:** Heinz Steffens, Alexander C. Mott, Siyuan Li, Waja Wegner, Pavel Švehla, Vanessa W. Y. Kan, Fred Wolf, Sabine Liebscher, Katrin I. Willig

## Abstract

Excitatory synapses on dendritic spines of pyramidal neurons are considered a central memory locus. To foster both continuous adaption as well as the storage of long-term information, spines need to be plastic and stable at the same time. Here we advanced *in vivo* STED nanoscopy to superresolve distinct features of dendritic spines (head size, neck length and width) in mouse neocortex for up to one month. While LTP-dependent changes predict highly correlated modifications of spine geometry, we find both, uncorrelated dynamics, as well as correlated changes, indicating multiple independent drivers of spine remodeling. The magnitude of this remodeling suggests substantial fluctuations in synaptic strength, and is exaggerated in a mouse model of neurodegeneration. Despite this high degree of volatility, all spine features also exhibit persistent components that are maintained over long periods of time. Thus, at the nanoscale, stable dendritic spines exhibit a delicate balance of stability and volatility.

## INTRODUCTION

Synapses on spines of principal neurons are a major locus of memory formation and maintenance in cortical circuits (*1–4*). To serve this function, spine synapses must be dynamic to change during learning, and simultaneously exhibit features of long-term persistence to maintain memory traces. Synaptic spines and their components are nano-physiological information processing devices (*5*) and the advent of superresolution now enables the assessment of their dynamics and the remodeling of their components at unprecedented resolution (*6–8*). Imaging in tissue imposes challenges to all superresolution light microscopy techniques and the first approach to overcome these in the mouse brain was STimulated Emission Depletion (STED) microscopy (*6*). So far, however, time periods relevant to long-term memory storage have remained unfeasible. In addition, previous approaches were technically elaborate and quite invasive, further limiting chronic long-term studies (*6, 9*). Refining the window implantation technique we here present an approach for long-term STED imaging of the cortex that enables high definition imaging without the need for image post-processing or expensive adaptive optics. With these advancements, we were able to monitor individual spines at superresolution for up to one month in the cortex of living mice, an order of magnitude longer than previous studies (*7*).

For the first time this allowed us to trace the dynamics of all features of spine geometry that can impact the long-term maintenance of synaptic strength. Consistent with the synaptic trace theory of memory formation, cortical engagement in learning tasks is often accompanied by a transient peak in spine generation, followed by the selective stabilization of newly formed spines (*10, 11*). In addition to plain spine turnover, the size of cortical spines can vary due to learning-induced and/or spontaneous processes (*3, 12, 13*), a process which affects all features of postsynaptic organization from receptor complement, postsynaptic scaffold (*14*) to the actin cytoskeleton, maintaining the spine’s morphology (*15–17*). Inducing synaptic long-term potentiation (LTP) *in vitro*, for instance, simultaneously leads to remodeling of the post-synaptic density (PSD) and to changes in spine morphology, including the concerted expansion in head size and neck width and shortening of neck length (*18–20*). Distinct from such activity-dependent changes in spine size, there is evidence arguing for ‘intrinsic’ spontaneous morphological volatility, independent of neuronal impulse activity and synaptic transmission, endowing the system with a large level of flexibility (*12, 13*). The contribution of both processes to structural remodeling under baseline conditions *in vivo*, however, remains poorly understood to date. If activity-dependent mechanisms in fact underlie the bulk of ongoing *in vivo* spine remodeling, one would expect that ongoing changes in spine head size, neck length and neck width are effectively controlled by a single underlying master process and therefore are tightly correlated, such as observed for LTP (*20*). Such concerted changes would optimally orchestrate the contributions of spine geometry changes to synaptic potentiation, since synaptic strength is predicted to substantially increase by shortening and widening of the spine neck (*21*). Remodeling of the spine as well as of the PSD is dependent on the actin cytoskeleton and is driven by postsynaptic Ca^2+^ influx (*17*). Because the actin cytoskeleton is composed of several pools of f-actin, all of which undergo continuous assembly and disassembly, not only the spine head but the entire spine morphology can be expected to exhibit spontaneous intrinsic fluctuations. However, while actin dynamics are well studied in the spine head, little is known about its dynamics in the spine neck. Morphologically, branched filaments dominate in the spine head whereas linear filaments are salient in the spine neck (*22*). Depending on whether and how alterations of head- and neck geometry are coordinated, such changes in total spine geometry may either enhance or suppress fluctuations of synaptic strength. In the past, studies on synaptic plasticity exclusively used the spine total fluorescence as a proxy of head volume and thus synaptic strength owing to limitations of optical resolution (*23–26*). As examining the dynamics of individual spine features requires long-term monitoring at nanoscale resolution, however, the questions of whether activity-driven or spontaneous remodeling is dominant *in vivo*, of whether there are one or many drivers of spine geometry remodeling and whether such drivers are independent or controlled by a single master process, remain unanswered to date.

Our data using long-term *in vivo* STED microscopy demonstrate that stable spines in fact undergo strong morphological fluctuations in all their geometrical features, even under baseline conditions. We show that the distribution of spine head sizes and neck width approximately is log-normal, indicating multiplicative dynamics, while neck length does not. Importantly, we discovered that the dynamics of some spine geometry features, such as neck length and head size, were uncorrelated, while changes in head size and neck width are correlated as expected for a process driven by activity-dependent synaptic plasticity. Although this remodeling is comparable in magnitude to activity-dependent processes, as observed after LTP-induction, and thus suggest substantial fluctuations in synaptic strength, changes *in vivo* might be regulated differently. Despite a high level of volatility, all spine features influencing synaptic strength also exhibit persistent components that are maintained over long periods of time. We furthermore find that the fate of spines to some degree can be predicted from their morphology, such that e.g. the spine head size of transient spines is on average only around 1/3 of that of stable spines. In addition, we investigated these morphological features and their dynamics in a mouse model of neurodegeneration, namely a transgenic model of Amyotrophic lateral sclerosis (ALS). This disease is characterized by the degeneration of upper and lower motor neurons. Chronic nanoscale imaging in the affected motor cortex of these mice revealed a pronounced loss of spines, paralleled by a highly dynamic increase in head size of the remaining stable spines, arguing for a large degree of synaptic remodeling of the remaining, persistent spines.

## RESULTS

### Chronic window implant for *in vivo* STED microscopy

In order to achieve nanoscale resolution of structural correlates of synapses over extended periods of time *in vivo*, we built a custom-designed STED microscope (Figure 1A). Our microscope consists of an upright microscope stand to which we attach a blue excitation (one-photon) laser to excite EGFP and an orange laser for stimulated emission depletion. A vortex phase plate in the STED-laser-beam creates a doughnut-shaped focal intensity pattern for superresolution in the xy-plane. Epi-fluorescence is detected via a confocal pinhole and single photon detector. The achieved superresolution during chronic *in vivo* imaging was around 96nm (Figure S1), which is 5–10 times higher than that of a conventional two-photon microscope. *In vivo* STED requires a mechanically stable and thermally isolated microscope to avoid thermal drift. To this end, we designed a mounting plate with a large heat sink (*8*). The cranial window needs to be of highest quality, since small optical aberrations can massively deteriorate image quality. Most importantly, the craniotomy needs to be as atraumatic as possible. In particular, we tested several parameters of the implantation procedure, aiming at a persistent short or negligible distance between cover slip and brain surface. This is crucial in order to minimize optical aberrations on the one hand and to avoid motion artefacts, associated with brain vessel pulsation on the other hand. We compared two different sizes of cover-slips (4 and 5 mm diameter). We observed less regrowth and thus superior image quality using a 4mm cover glass. Due to the curvature of the skull the 4mm coverslip was easier to fit into the craniotomy directly in contact with the brain surface. Another important factor was the dental cement. We tested a two-component, UV-light curable dental cement (Paladur^®^) and a self-curing adhesive resin cement (SuperBond C&B^®^). In our hands, SuperBond C&B^®^ was superior to the Paladur^®^ cement and resulted in an improved window quality, with less motion-related artefacts. To affix the mouse’s head underneath the objective, we designed a novel head bar (Figure S1A), which can be cemented flush to the skull to allow access to a high numerical aperture (NA, 1.3), short working-distance objective. To protect the surface of the window from scratches and dirt, we applied a layer of protective silicone on top of the window at the time of the window implantation and between the imaging sessions. The silicone can easily be removed before imaging and without the need to further clean the window.

**Figure 1:**
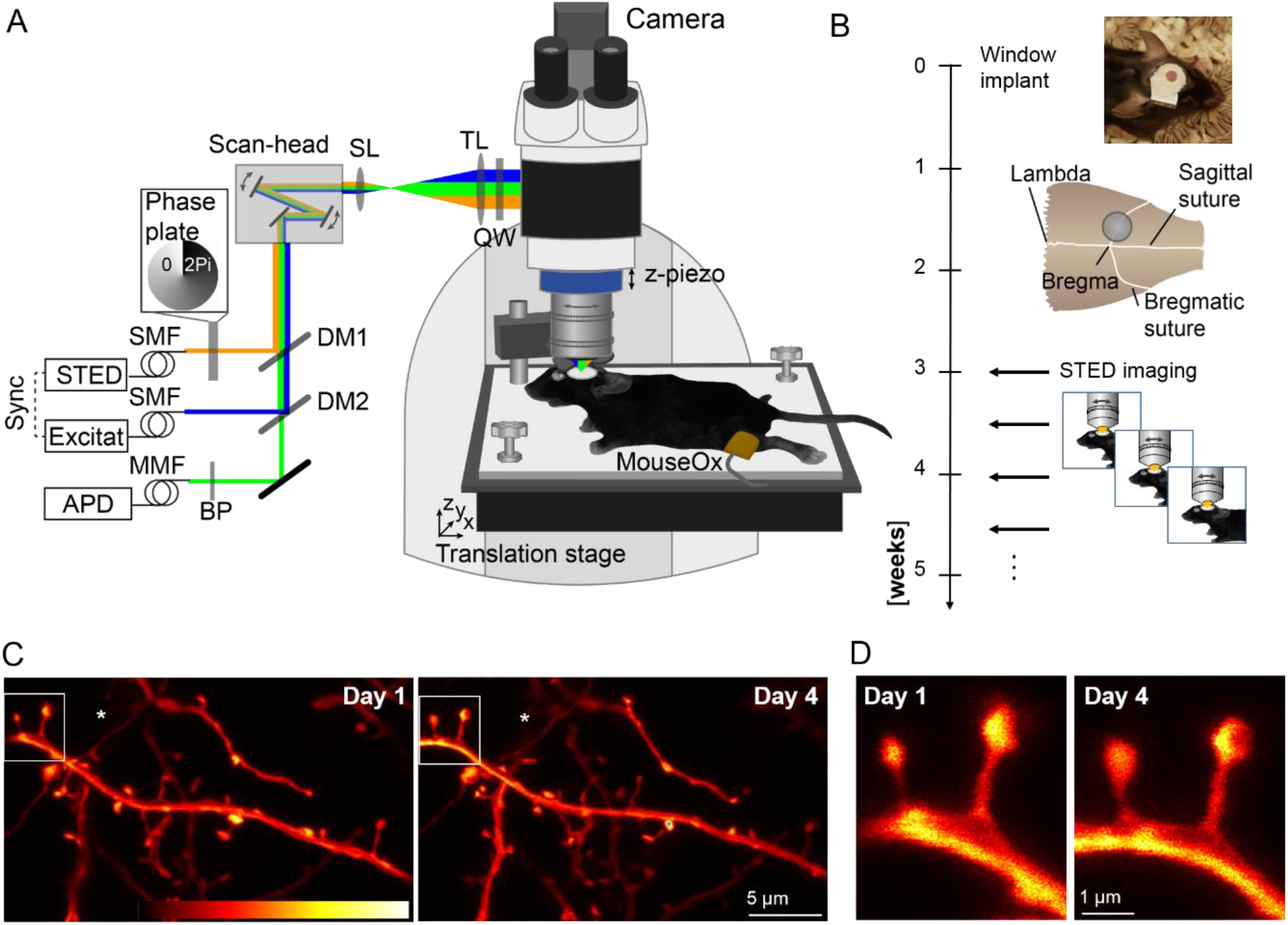
Repetitive superresolution of the mouse motor cortex using STED microscopy. (A) Microscope design: A custom-made STED microscope is attached to a microscope stand. The pulsed 483 nm excitation is temporally synchronised electronically with the 595 nm STED light pulses and merged spatially by dichroic mirrors (DM). After passing two galvanic mirrors in the scan-head the light is imaged by a scan (SL) and tube lens (TL) before being focused by a glycerol immersion objective (numerical aperture 1.3) with a correction collar. The mouse is mounted via a head bar on an adjustable heating plate. Vital functions are controlled by a pulse oximeter (MouseOx). (B) After window implantation and a three week recovery period the mouse was imaged twice a week. (C) Representative raw data example of an apical dendrite of a pyramidal neuron in motor cortex of a Thy1-GFP-M mouse imaged at day 1 (left) and day 4 (right). An axon captured in the same field of view is marked by (*). (D) Magnification of marked region in (C). Images are maximum intensity projections of 6 frames. Abbreviations: APD: Avalanche photo diode detector, BP: band-pass filter, MMF: Multi-mode fibre, QW: Quarter wave plate, SMF: Single-mode fibre. Colorbar: 0-212 photon counts.

### Longitudinal superresolution STED microscopy

*In vivo* STED imaging commenced after a recovery period of 3-4 weeks (Figure 1B). The same field of views were revisited twice a week. Mice were anaesthetised, using a combination of Midazolam, Metedomidin and Fentanyl. The mounting plate is tiltable and the cranial window was accurately aligned perpendicular to the optical axis of the microscope (*8*). We frequently observed sprouting of fine new blood vessels underneath the coverslip, which can affect image quality (Figure S1B). STED microscopy in the motor cortex of a Thy1-GFP-M transgenic mouse (GFP-M line) (*27*) yielded crisp images depicting dendritic spine morphology in layer 1 at nanoscale resolution (Figure 1C,D). The same dendrite was imaged three days later, showing changes in spine morphology (Figure 1C,D). STED image resolution critically hinges on the spectroscopic properties of the fluorescent molecule and on the STED focal doughnut. To correct for spherical aberrations, we adapted the correction collar of the objective at each field of view. To determine the resolution most accurately, we measured the full-width at half-maximum (FWHM) within the *in vivo* images. The smallest/thinnest structures were axons (*in Figure 1C, Figure S1C), which could be resolved at a FWHM of 96nm (Figure S1C), an upper estimate of the resolution. With these settings we recorded STED microscopy images over a period of up to 28 days (Figure 2), which enabled us to monitor fine changes in spine morphology, such as the spine neck (Figure 2, insets) that are typically obscured when using e.g. two-photon microscopy. In addition, these imaging data also enabled us to detect individual dendritic spines of all sizes with high precision and to observe morphological phenomena such as clustered spine formation (Figure S2A) or ‘touching heads’ (Figure S2B).

**Figure 2:**
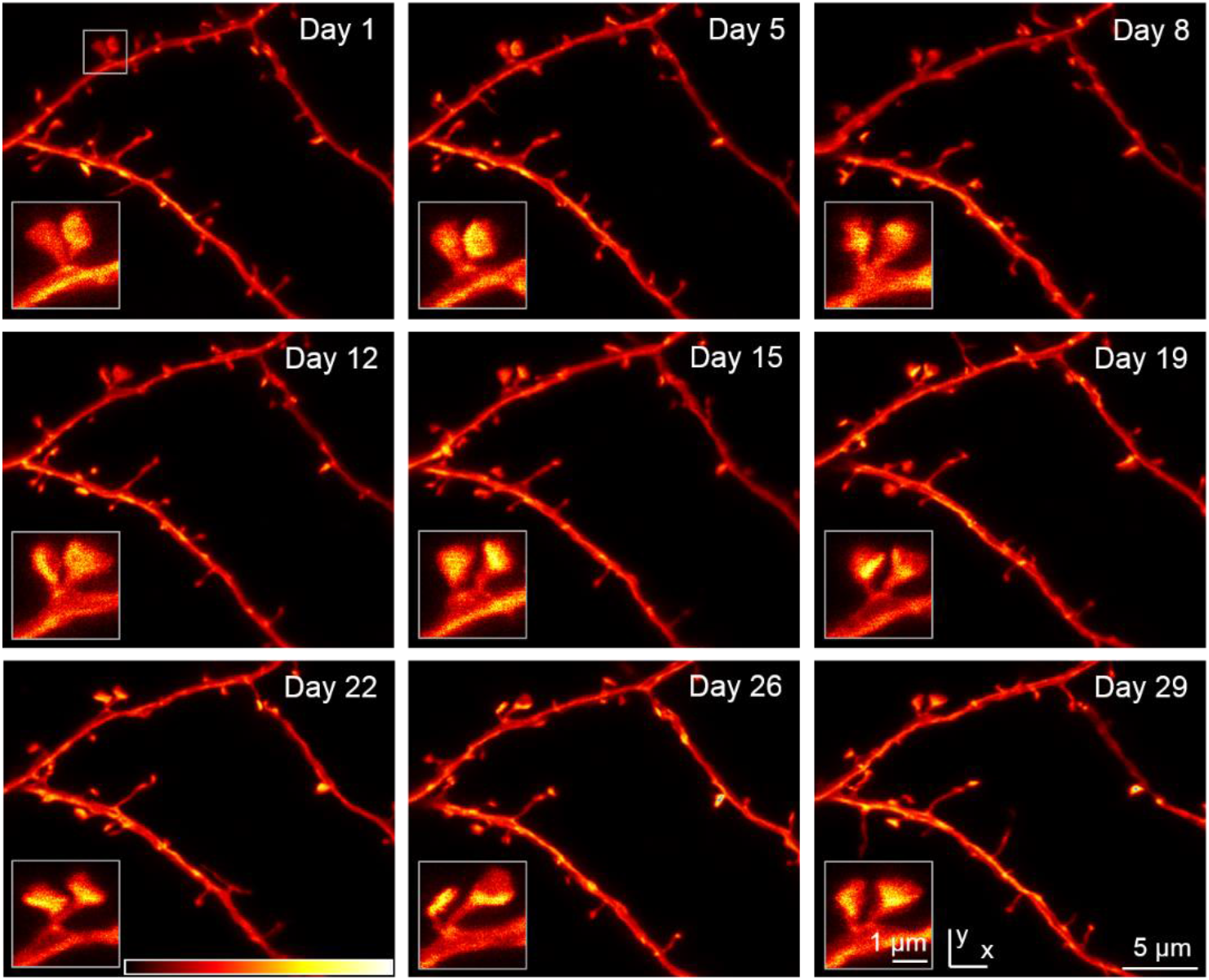
Chronic STED imaging of dendritic stretches in layer 1 of motor cortex. Superresolution reveals changes of spine nanoplasticity of large, mushroom-type, stable spines (inset). Images are maximum intensity projections raw data. Colorbar: 0-120 photon counts.

### Distribution and interdependency of spine parameters

Dendritic spines, emanating laterally from the dendrite within the same or a consecutive focal plane to the dendrite were analysed across all time points (Figure 3A). The longest axis of the head (head length) and the axis perpendicular across the spine head (head width) were measured. The size of the head cross section was approximated by computing the area of an ellipse from these parameters. Neck length was defined as the distance from the base of the spine neck to the beginning of the spine head (Figure 3A). The neck width represents the thinnest extent of the spine neck (Figure 3A). We observed a large range of head sizes (median: 0.31μm^2^, interquartile range (IR): 0.23–0.43μm^2^) and neck lengths (median: 803nm, IR: 485–1154nm), spanning in total an order of magnitude (Figure S3A,B), while the neck width (median: 238nm, IR: 208–273nm) was less variable (Figure S3C). All three parameters are positively skewed and therefore we plotted the histogram of the log_10_ values for the three prime parameters (Figure 3B-D). Interestingly, we observed a log-normal distribution for head size and neck width, but not for neck length.

**Figure 3:**
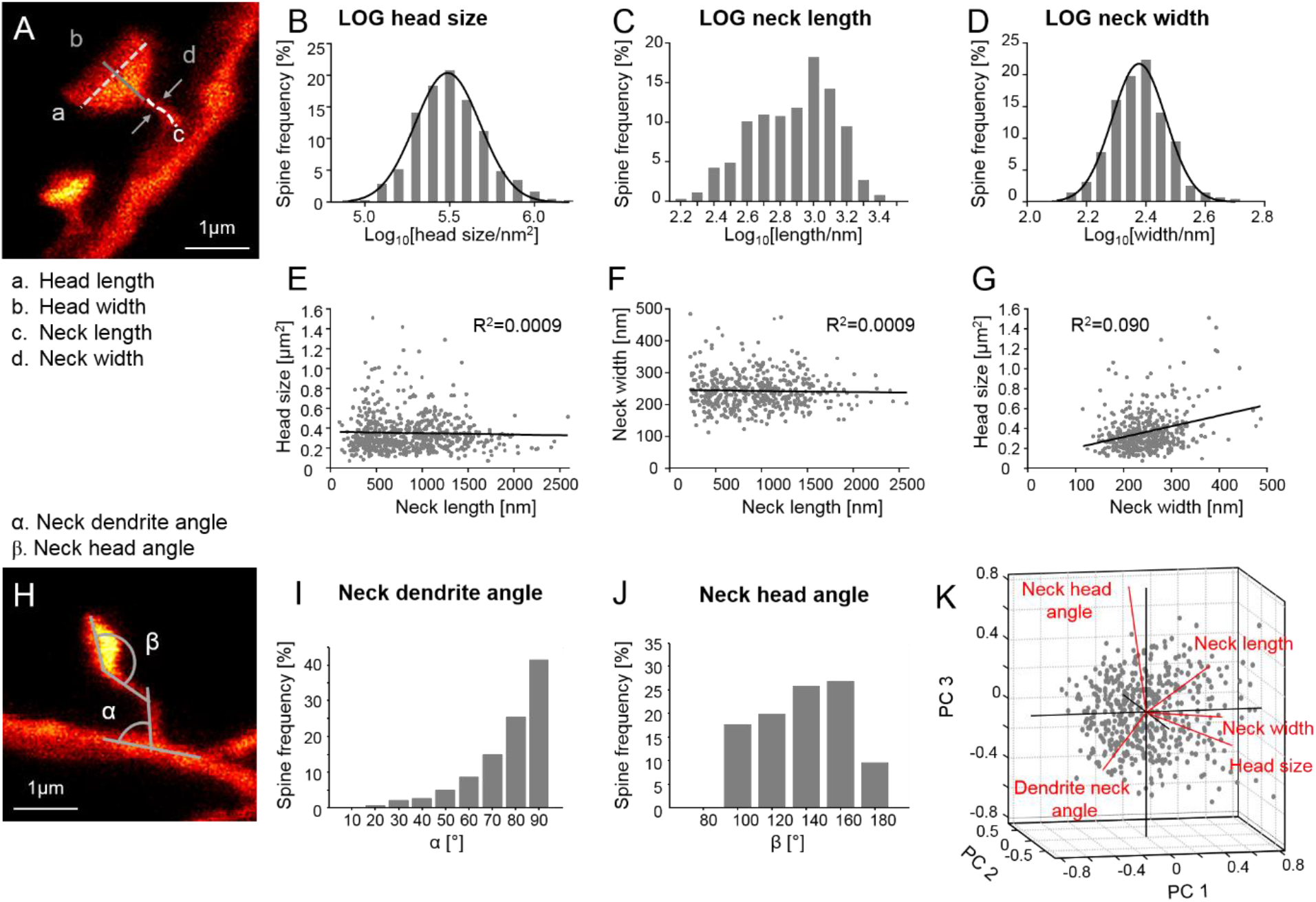
Spine morphometric parameters are largely uncorrelated while head size and neck width but not neck length exhibit multiplicative dynamics. (A) Features of stable spines assessed are head length, head width, spine neck length and neck width. (B-D) Histogram of the logarithmic spine head sizes (B), spine neck lengths (C) and neck width (D). (B+D) The Log_10_ data is normally distributed indicated by a Gaussian fit (black line). (E–G) Correlation between spine parameters. Spine head size (E) and neck width (F) as a function of neck length and head size as function of neck width (G); linear regression (black). (H–J) The angle between the dendrite and the spine neck (α, neck dendrite angle) (I) and between the neck and the spine head (β, neck head angle) (J) is measured. (K) Principal component (PC) analysis of 5 morphological parameters.

To assess whether the morphological parameters were interdependent, we investigated their correlation. The parameter neck length did neither correlate with head size (R^2^<0.001, p=0.46, Figure 3E) nor with neck width (R^2^<0.001, p=0.52, Figure 3F); meaning that, e.g. large spine heads are attached to short and long necks with equal probability and that the width of a spine neck is independent of its length. A weak, but highly significant, positive correlation, however, was observed for the parameters head size and neck width (R^2^=0.09, p<0.0001, Figure 3G). We then measured the angle between the spine neck and dendrite (neck-dendrite angle) as well as the angle between the base of the spine head and the neck (neck-head angle; Figure 3H–J). The majority of spines extended from the dendrite at an angle of ~90°. A principal component analysis (PCA) of the five morphological parameters (head size, neck width/length, neck-dendrite angle and neck-head angle) of both stable and transient spines revealed that the parameters head size, neck length and neck width contribute to the first component and thus that those parameters most strongly determine the variability within the data set (Figure 3K). Moreover, the eigenvalues (1.48, 1.05, 1.0, 0.87, 0.6) of the z-scored data were rather similar and therefore all morphological parameters are largely independent. In other words, we have no evidence for a clear interdependency of the morphological parameters we measured. Besides spines bearing a defined spine head, we could also superresolve filopodia (long protrusions lacking an obvious head), which we analysed separately (Figure S3D-H). Filopodia were on average 2300nm (±762nm SD) long (Figure S3E), with a neck width of 236nm (± 49nm SD, Figure S3F) and emanated from the dendrite at an angle of 75.8° (64.9–79°, 95% CI) (Figure S3G). All filopodia in our data set occurred only once, thus had a lifetime of less than 3 days (likely rather minutes to hours). We did not find a correlation between the width and length of filopodia (R^2^=0.14, Figure S3H).

### Temporal changes of stable spine morphology

We next asked how those morphological parameters change over time. To this end, we analysed head size, neck length and neck width for stable spines over time (Figure 4A–I). On average all measures stayed stable across all time points indicating the absence of phototoxic effects (Neck length p=0.85, head size p=0.55, neck width p=0.1; Kruskal-Wallis and Dunn’s multiple comparison test; Figure 4B). The majority of spines underwent relative changes exceeding ±10% of the initial spine head size or neck length over a period of 3–4 days (Figure 4C). More specifically, only 22% of spines displayed a minor head size change within ±10%. However, 44% of spines decreased in head size more than 10%, while 34% increased in head size exceeding 10%. 25% of spines underwent a neck length variation within ±10%, while 39% of spines decreased in neck length and 36.3% of spines increased in neck length exceeding 10% (Figure 4C). On average, spines grew in head size by 26% (median, IR: 11–47%) (Figure S4A,B) and neck length by 22% (median, IR: 9–45%) (Figure S4C,D), up to a maximum of >200%. The median shrinkage of head size was - 21% (IR: −12 to −35%)(Figure S4B) and of neck length −23% (IR: −11 to −34%)(Figure S4D). Furthermore, we analysed for each spine parameter the changes as a function of their initial size. The changes in head size and (Figure 4E) and neck length (Figure 4F) were larger for larger initial size and were overall negatively correlated to the initial size for head size (R^2^=0.097, p<0.0001; Figure 4E) and neck length (R^2^=0.078, p<0.0001; Figure 4F), indicating a preference for shrinkage of large spines and growth of small spines. The same tendency is observed when plotting the percentage change of head size and neck length (Figure S4A,C). Normalized relative changes of the head size were not correlated with normalized relative changes of the neck length (Figure 4D). Please note that the normalization restricts the changes to +1 and −1; the distribution is therefore symmetric.

**Figure 4:**
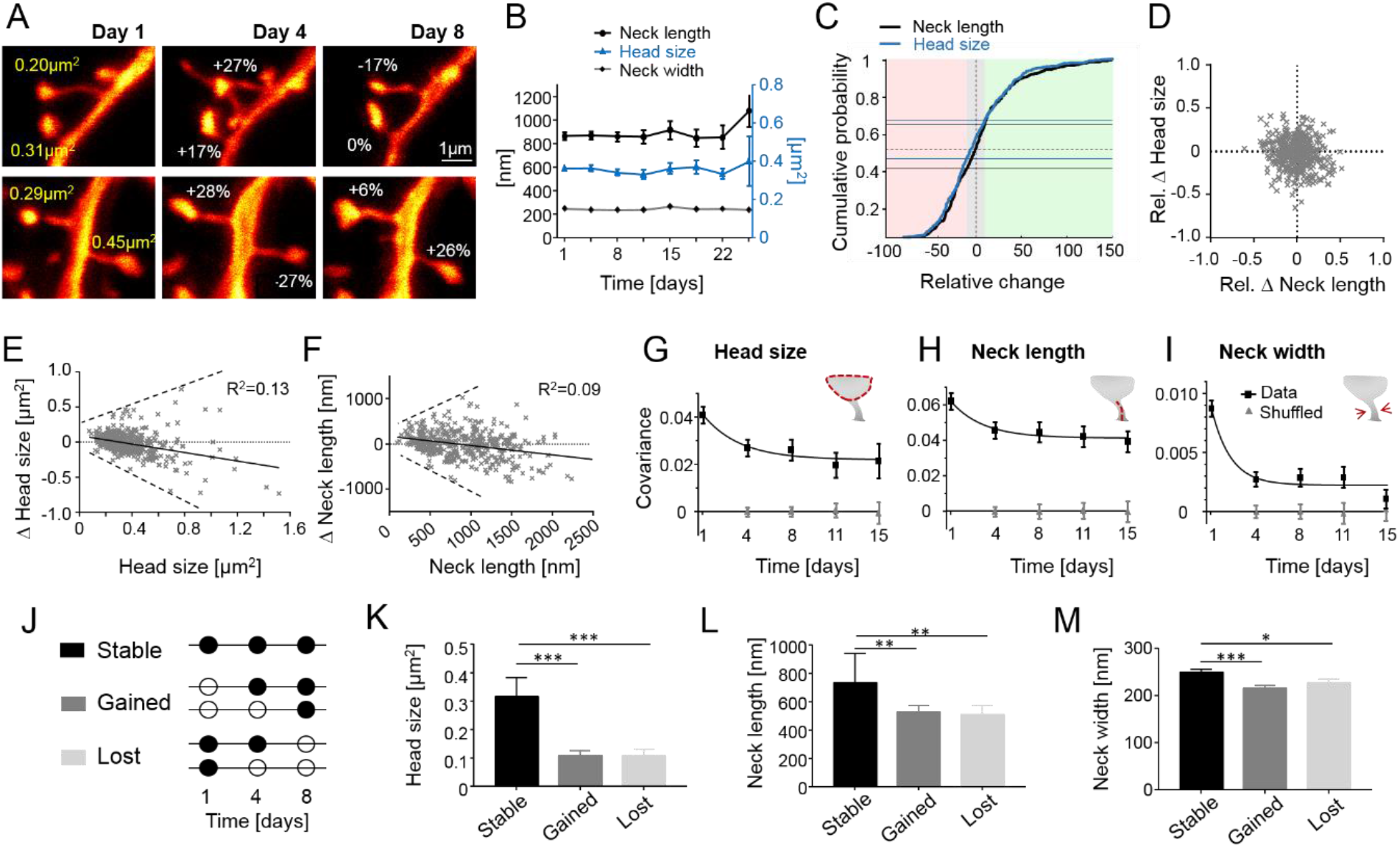
In persistent spines head size and neck length fluctuate independently and are more persistent than fluctuations in neck width, while in transient spines all spine parameters predict gain and loss. (A) Representative examples of changes in spine head size. (B) The morphological parameters are stable over the observation period of 24 days. (C) Cumulative distribution of relative changes of neck length (black) and spine head size (blue) over 3–4 days. Fraction of spines that changed spine head size within ±10% (gray area) is indicated by blue horizontal lines, while the same for neck length is indicated by black horizontal lines (light red area denotes relative change in size to lower than −10% while changes exceeding 10% are indicated in green area). (D) Normalized changes of head size and neck width are not correlated. (E) Absolute changes in head size over 3–4 days increase with initial size (dashed line is a guide to the eye) and are negatively correlated. (F) The absolute change in neck length over 3–4 days increases with initial neck length (dashed line is a guide to the eye) and are negatively correlated. (G–I) Covariance of morphological spine parameters (logarithmic values) across imaging sessions and mono-exponential fit. Error bars are SD of bootstrapped data. (J) Morphological analysis depending on spine history (open circle – spine not present; filled circle – spine present). (K–M) Gained and lost spines show significantly smaller head sizes (K) and smaller neck length (L) and thinner neck width (M). *p<0.05; **p<0.01; ***p<0.001. Data are mean ±SEM (B,M) and median +95% CI (K,L).

While these data capture the changes between two consecutive time points, we wondered how the morphology changed over longer imaging intervals. Do spines continue to grow or shrink? To address that question, we computed the covariance function for up to 15 days for the variables head size (Figure 4G), neck length (Figure 4H) and neck width (Figure 4I). All three parameters showed a drop in the covariance function after the first time interval of 4 days and then levelled out with a large offset for head size and neck length and low offset for the neck width. The interpretation of these analyses is illustrated for a model of temporal changes of a spines of different head sizes in Figure S5A-C. For a multiplicative dynamics, head sizes are log-normally distributed and the temporal changes in size (□M) on a log-scale have the same standard deviation. On a linear axis, the standard deviation would increase in size with average spine size as expected for multiplicative changes (Figure S5A, inset). While the covariance function for temporal fluctuations of an individual spine decays to zero (Figure S5B) the covariance function of the whole ensemble exhibits an offset (Figure S5C) that is equal to the variance of the distribution of sizes across the population. This means that the presence of such an offset in our measurement (Figure 4G–I) indicates that large heads mainly remain large and small heads remain small over the imaging period of 15 days. Therefore, the main fluctuations in size occur at time scales of 3–4 days or shorter and are largest for neck width. For comparison between the parameters, we plotted also the auto-correlation, i.e. the normalized covariance function (Figure S5D). The largest drop in auto-correlation was observed for the neck width. The correlation between our parameters, i.e. the cross-correlation was close to zero between head size and neck length as well as between neck length and width (Figure S5D). However, a 20–30% correlation for time intervals up to 15 days was observed between head size and neck width.

### Gained and lost spines have smaller heads than stable spines

Next, we asked whether the changes in spine morphology are related to the spine fate or previous history. We analysed all spines for 3 consecutive time points and categorized them into stable, gained and lost spines based on their lifetime (Figure 4J). As predicted, stable spines (present on all three imaging time points) had an almost threefold larger head compared to spines just gained (gained) or spines measured at the time point prior to their loss (lost) (head size stable spines 0.32 (0.26–0.38) μm^2^, gained 0.11 (0.09–0.13) μm^2^ and lost spines 0.11 (0.09-0.13) μm^2^ (median and 95% CI; stable vs gained p<0.0001; stable vs lost p<0.0001; Kruskal-Wallis test with Dunn’s multiple comparison test; Figure 4K). Similarly, stable spines had a longer neck (neck length stable spines 737 (642–945) nm, gained 531 (437–573) nm and lost spines 511 (437–576) nm (median and 95% CI; stable vs gained p = 0.0019; stable vs lost p=0.0016, Kruskal-Wallis test with Dunn’s multiple comparisons test; Figure 4L) and a larger neck width (neck width stable spines 249.4 ±6nm, gained 215 ±7nm and lost spines 228 ±8nm compared to gained and lost spines (mean ±SEM; stable vs gained p=0.0006, stable vs lost p=0.032; ANOVA with Dunnett’s multiple comparison test; Figure 4M).

### Probing ultrastructural morphological abnormalities of dendritic spines in a transgenic mouse model of ALS

Finally, we applied chronic STED microscopy to investigate morphological alterations of dendritic spines in motor cortex of a transgenic mouse model of Amyotrophic lateral sclerosis (ALS). ALS is a fatal disease primarily caused by the degeneration of upper- and lower motor neurons in motor cortex and spinal cord, respectively (*28, 29*). We employed a well-characterized mouse model of the disease that is based on the overexpression of the mutated Superoxide-dismutase-1 gene (SOD1^G93A^, hereafter called SOD (*30*)). These mice recapitulate key phenotypic features of ALS and die prematurely due to paralysis. Earlier work indicates that upper motor neurons, which reside in cortical layer V, are also affected in the mouse model but actual insight into ultrastructural abnormalities and the dynamics of those changes *in vivo* is lacking to date (*31–33*). In order to investigate dendritic spines of layer V pyramidal neurons in SOD1 transgenic (tg) mice, we crossed SOD1^G93A^ with GFP-M mice and examined both the SOD1^G93A^ expressing mice as well as their non-transgenic littermates (WT). First, we assessed the spine density of apical tufts of layer V pyramidal neurons over three consecutive time points (Figure 5A) and found a significant decrease in spine density in SOD tg mice (effect of group: *F*_1,56_=14.34, *p*=0.0007; effect of time: *F*_2,56_=2, *p*=0.15; group-by-time interaction effect: *F*_2,56_=1.39, *p*=0.26, two way repeated measures ANOVA; Figure 5B). Overall, spine density stayed stable within the 8 day imaging period. A detailed morphological assessment of stable spines revealed that the distribution of the parameters head size, neck length and neck width in the SOD mouse was similar to those found in WT mouse (Figure S6A–C). We observed a log-normal distribution only for head size and neck width, while the neck length was not log-normally distributed (Figure S6A–C). On average, spine head size (median: 0.35μm^2^, IR: 0.24–0.47μm^2^) was increased and showed a larger variance in SOD mice (head size: p=0.011, two-sided, unpaired t-test with Welch’s correction; head size variance: p<0.006; F-Test; Figure 5C). The neck length (median: 824nm, IR: 541–1110nm), on the other hand, did not differ significantly between genotypes (neck length: p=0.62, two-sided, unpaired t-test with Welch’s correction; neck length variance: p=0.27; F-Test; Figure 5D). The neck width (median: 229nm, IR: 200–264nm) showed a small, but significant, decrease in average size for SOD mice, but not in variance (Figure S6D). We also compared the relative changes of head size and neck length between two consecutive imaging sessions (Figure 5E) and also observed an increase in the variance of relative head size changes (p=0.008, two-sided, unpaired t-test with Welch’s correction), but no difference in the variance of neck length changes (p=0.47, two-sided, unpaired t-test with Welch’s correction). In summary, the spine density in SOD mice is decreased, while the heads of the remaining spines are larger, have a greater variance and undergo a more pronounced change in size over time. Neck length did not differ between WT and SOD mice, neither in absolute values nor in variance or in the magnitude of their temporal change.

**Figure 5:**
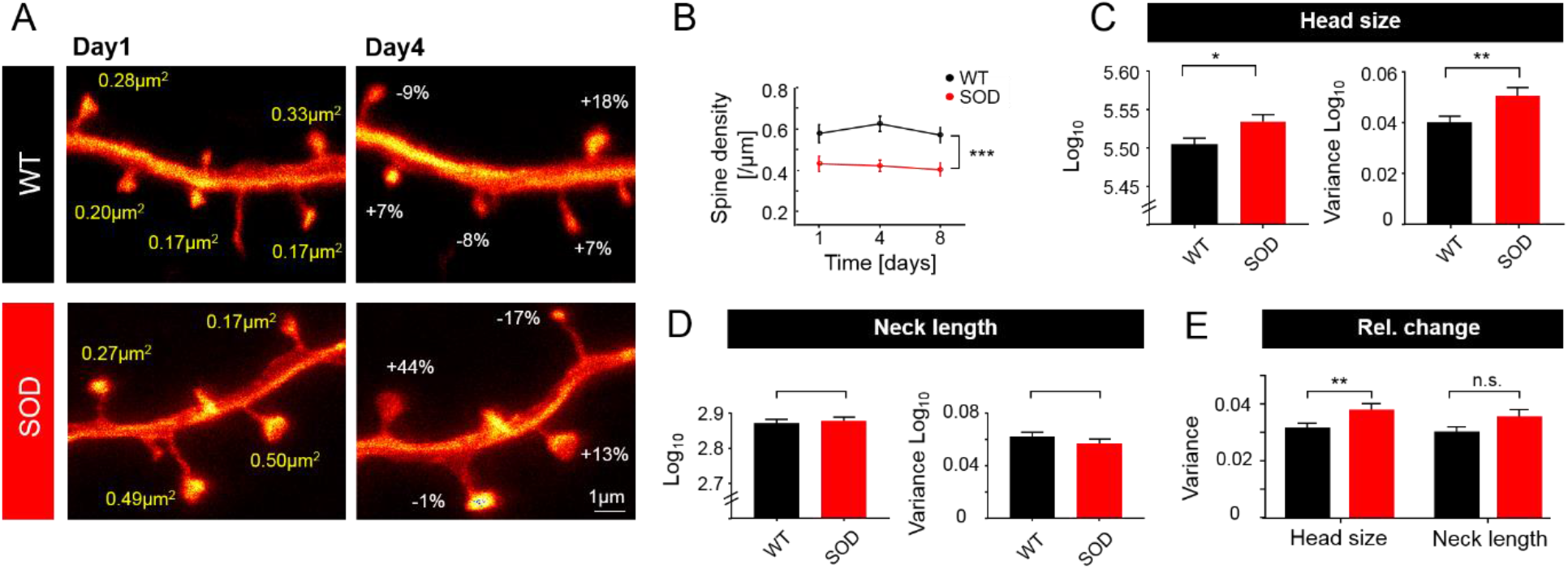
Spine density in SOD transgenic mice is reduced, while fluctuations in remaining spine heads are exaggerated. (A) Representative example of a GFP-expressing dendrite in a WT and a SOD transgenic mouse showing changes in spine head size within 3 days (images are maximum intensity projections). (B) WT mice show higher spine density than SOD mice. (C) Head size of stable spines as well as their variance were increased in SOD mice. (D) Lack of difference in neck length or variance of neck length in SOD mice. (E) Variance of relative changes over 3–4 days of spine head sizes is significantly increased in SOD mice, while the increase in neck length is not significant. *p<0.05; **p<0.01; ***p<0.001. Data in (B–E) are mean ±SEM.

## DISCUSSION

Establishing chronic *in vivo* STED nanoscopy to superresolve dendritic spines in mouse neocortex for up to one month enabled us to provide the first characterization of ongoing dynamic fluctuations in spine head and neck geometry at nanoscale resolution over long periods of time *in vivo*. We found that all assessed geometric features exhibit spontaneous fluctuations of substantial magnitude. Ongoing changes in spine geometry are of a magnitude similar to changes caused by LTP induction and they are presumably indicative of substantial modifications of synaptic strength. Our data for persistent spines are consistent with the assumption that their geometric features fluctuate around mean values that are spine specific and maintained over periods on the order of months at least. The maintained components of spine head size and spine neck width were matched such that larger spine heads are systematically associated with wider spine necks despite substantial ongoing fluctuations. For neck length and head size, temporal fluctuations around their respective mean values were statistically independent of each other, indicating the existence of multiple independent drivers of geometric remodeling. Confirming the predictions of computational models for the random cooperative molecular assembly and turnover of postsynaptic supra-molecular complexes, spine neck width and head size exhibited approximately log-normal distributions and multiplicative dynamics. The observed volatility of synaptic spine morphology was exaggerated in a mouse model of neurodegeneration. Together these findings provide a picture of *in vivo* spine dynamics exhibiting a delicate balance of stability and volatility at the nanoscale level.

### Longitudinal STED nanoscopy to characterize dendritic spine nanostructure

We here report the development of chronic *in vivo* STED imaging of synaptic structures in the neocortex for up to one month. Key to the procedure is a chronic stable cranial window preparation, with minimal amounts of fluid in between the coverslip and the cortical surface as well as minimal regrowth of adhesive tissue/bone or dural thickening. We focused our method on decreasing the distance between cover slip and brain surface, since motion artefacts were always present as soon as this distance exceeded 10–20μm. The key development of the method was an atraumatic craniotomy, the right size of the cover slip, resin cement and the design of a novel head bar, allowing access to a high NA objective. This optimized protocol enabled us to successfully scan 100% of the mice implanted, chronically over at least three weeks recording 2 to 9 time points per field of view.

We assessed five distinct spine parameters: size of the spine head cross section, length of the spine neck, neck width and the spine neck-head and spine neck-dendrite angles. The values we obtained *in vivo* are well in line with electron microscopy (EM) analyses of fixed tissue. For instance, the average neck diameter in our data was 238nm and the median head size was 0.31μm^2^, corresponding to a spine head volume of 0.13μm^3^ (assuming a spherical volume). These values are consistent with EM data (*34*). However, in sharp contrast to EM, our STED approach allows for a longitudinal assessment of synaptic structures *in vivo*. The log-normal distributions we observe for head size and neck width are well explained by mathematical models based on multiplicative dynamics (*35, 36*). The right-skewed but not log-normally distributed neck length, however, neither follows multiplicative nor additive dynamics and does not fit to existing models.

### Stability and volatility of spine geometry of persistent spines

In this study, we have analysed temporal changes of dendritic spine geometry, to our best knowledge for the first time, chronically in cortex of living mice for up to one month. In contrast to most longitudinal *in vivo* studies, assessing structural plasticity by considering spines as binary entities, our nanoscopy approach enabled us to investigate dedicated spine parameters at ultrastructural resolution. It is well accepted that the size of a dendritic spine scales with the strength of the synapse and is predictive of its lifetime (*37, 38*). However, recent evidence also argues for a critical impact of other parameters, such as neck length or width in determining the function of a synapse (*18, 20, 21*). As such, earlier studies using two-photon imaging demonstrated that the neck length shortens with the potentiation of a spine (*18, 19*). Conducting STED imaging in brain slices combined with glutamate uncaging suggests that the most critical parameter determining dendritic spine compartmentalization is the spine neck (*20*). Moreover, a novel electro-diffusion model revealed the substantial impact of spine neck geometry on synaptic strength (*21*). Our data set, acquired *in vivo*, under baseline conditions (that is without the application of a dedicated learning/memory paradigm), shows that both spine heads and necks fluctuate significantly. We particularly focused our investigation on stable spines, which are believed to embody structural correlates of learning and memory (*2, 11*). The majority of those (~80%) underwent a fluctuation in head size and neck length of more than 10% (~40% even of more than 30%) within 3–4 days. While these alterations might seem small at first glance, they could readily affect the strength of the corresponding synapse. For comparison, synaptic potentiation triggered by chemical LTP stimulation or glutamate uncaging has been shown to cause an increase in spine head size between 20–40% in a large fraction of spines (*39–42*) as well as a change in neck length by ~20–30% (*18, 20*) and neck width increase by 30% (*20*). The interpretation of those data, however, is complicated by the fact that these studies are conducted *in vitro* and rely on artificial and probably highly potent stimulation protocols. Our data demonstrate that similar effect sizes occur *in vivo* already under baseline conditions, even in the absence of a dedicated stimulation protocol. Importantly we already observed such large changes within 3–4 days, but also witnessed a large offset of 50–70% in the auto-correlation of head size and spine length over 15 days, indicating overall size stability. This suggests that spine geometry might be volatile within rather short time frames of days, but able to maintain a mean value over larger periods of time. Moreover, the geometric parameters as well as their dynamics were largely independent of each other. A small, yet persistent positive correlation was only observed between neck width and head size. Our data thus indicate that *in vivo* morphological parameters of dendritic spines exhibit a subtle balance of volatility and persistence. Intriguingly, the nature of these dynamics are consistent with the view that *in vivo* changes are dominated by independent turnover of the spines’ f-actin pools and rejects the predominant idea that the spine cytoskeleton is a passive component downstream to and driven by the primary processes of synaptic plasticity.

While most of the time-lapse studies to date address changes in spine volume or spine head size, little was known about spine neck changes *in vitro* and *in vivo*. We observed that changes of spine head size and neck length increased with their initial size and were negatively correlated, indicating that small spines tend to increase whereas large spines tend to shrink. This relationship supports models based on multiplicative dynamics (*35, 36*). Interestingly, in contrast to the distribution of head sizes, neck lengths did not follow a log-normal distribution and might thus not obey multiplicative dynamics, again supporting the notion that individual spine features are controlled by distinct drivers. In the future a revised model of spine geometry should provide better insights in the role of the spine neck in synaptic plasticity (*21*).

### Transient spines differ morphologically from persistent spines

Spine morphology is indicative of synaptic strength and also of its lifetime, but insight into the actual dimensions/magnitude is still lacking. We found a striking difference in head size as persistent spines were almost three times larger in head size compared to transient spines. They also possessed a 35% longer and 25% thicker neck compared to transient spines.

The most extreme case of transient spines are filopodia, which lack an actual spine head. These structures are on average 2.3μm long and possess a neck width of 236nm. All identified filopodia in our data set only occurred once, thus have a lifetime shorter than 3–4 days (see also (*37*)). While we cannot fully exclude that newly formed spines initially underwent a stage reminiscent of a filopodium, our current data argues against a major role of filopodia acting as precursors of dendritic spines in adult mice under baseline conditions. Most newly formed spines were in fact much shorter than filopodia and stable, mature spines. Future studies are thus needed to explore the relevance of those immature structures.

### Ultrastructural alterations of dendritic spines in SOD transgenic mice

We also performed nanoscopy in a transgenic mouse model of ALS. Prior to frank neuronal degeneration and loss, motor neurons are likely already impaired over a prolonged time in this disease. However, little is known to date about how neurodegenerative processes in ALS affect dendritic spines. We here monitored spines of layer V pyramidal neurons in a well-characterized model of ALS that is based on the overexpression of the mutated Superoxid-Dismutase-1 gene (SOD1). We assessed the density and dynamics of morphological parameters of dendritic spines during the presymptomatic stage (age 5-6 months) of low copy SOD1tg mice (*43*). We observed a pronounced decrease in spine density, which remained stable throughout the imaging period. Our data corroborate earlier studies conducted in a more aggressive mouse model (higher copy number (*31, 32*)) using Golgi Cox staining. The remaining stable spines in SOD mice had on average larger spine heads and wider necks compared to WT mice. Furthermore, head sizes varied more strongly not only at any given time point but also their changes over time were more variable compared to WT mice. The head size increase we observed could well represent homeostatic scaling to counteract for the loss of synaptic input (*44*) as a recent study demonstrated an increase in synaptic size and a broader distribution of the same in silenced neuronal networks (*45*). The enhanced dynamics of spine head sizes in SOD tg mice moreover argue for a higher level of synaptic remodeling, hence synaptic instability. Collectively, these results argue for structural modifications of upper motor neurons, which precede overt neuronal loss and symptom onset and substantiate the notion of ALS being a synaptopathy (*46*).

Taken together, we here for the first time established long-term *in vivo* STED microscopy in cortex of mice. Our data demonstrate that hitherto believed stable dendritic spines undergo pronounced morphological changes. Individual morphological features are largely independent, suggesting diverse drivers of synaptic plasticity.

## METHODS

### Animals

All mouse experiments were performed according to the guidelines of the national law (Tierschutzgesetz der Bundesrepublik Deutschland, TierSchG) regarding animal protection procedures and approved by the responsible authorities, the Niedersächsisches Landesamt für Verbraucherschutz und Lebensmittelsicherheit (LAVES, AZ 33.19-42502-04-17/2479) and Regierung von Oberbayern (AZ 55.2-1-54-2532-11-2016). Thy1-GFP (M-line, hereafter called WT) (*27*) and crosses of SOD1^G93A^ (*30*) and GFP-M mice (referred to as SOD) were used in groups of up to 5 mice per cage, with ad libitum access to food and water. Mice were kept at a 12/12 hour light/dark cycle. Mice were implanted at the age of 19 – 23 weeks and imaging commenced at 23-27 weeks of age. In total 7 males and 4 females, 5 WT (2m and 3f), 6 SOD (5m, 1f) were used.

### Mouse surgical procedure

The mouse was anaesthetised by intraperitoneal injection of a mixture of Fentanyl (0.05mg/kg), Midazolam (5mg/kg) and Medetomidine (0.5mg/kg). Once anaesthetised, the mouse was placed on a heating plate and shaved on the hind leg, as well as the surgical area on the scalp. During surgery and *in vivo* imaging vital functions and depth of anaesthesia were controlled: the body temperature was monitored with a rectal temperature probe, O2 saturation of the blood and heart rate were monitored using a pulse-oximeter (MouseOx STARR^®^, STARR Life Science Corp., Oakmont, PA) placed on the shaved thigh. A mixture of 50 vol% N2, 47.5 vol% O2 and 2.5 vol% CO2 was administered over a cone in front of the mouse’s nose to keep the oxygen saturation at ~98%. The mouse was positioned in a stereotaxic frame (Narishige, Tokyo, Japan) and the fur above the skull was cut. After sealing the edges of the skin with the tissue adhesive n-Butyl cyanoacrylate (Histoacryl^®^, B. Braun Melsungen AG, Melsungen, Germany) the skull was cleaned using a micro curette (10082-15; Fine Science Tools GmbH, Heidelberg, Germany) or drill. Next, the head bar was fixed with dental cement (Super-Bond^®^ C&B, Sun Medical Co. LTD, Japan) to the skull. After hardening of the cement, the mouse was moved to an adjustable and heated mounting plate with the head bar screwed to the head holder (Figure 1A, S1A). A circular craniotomy (4 mm diameter) was then performed (Drill: 216804; RUDOLF FLUME Technik GmbH, Essen, Germany; drilling head: HP 310 104 001 001 007; Hager & Meisinger GmbH, Neuss, Germany) centred over the motor cortex. The dura was gently removed with a fine biology tipped forceps (Dumont #5 biology, Fine Science Tools GmbH, Heidelberg, Germany). Care was taken to not damage the cortical surface and to avoid blood cell deposits at the region of interest. The cover glass of 4 mm diameter (Warner Instruments, CT, USA) was fit in tightly into the opening and affixed to the skull using tissue adhesive. Once held in place, the window was firmly fixed using dental cement. The surface of the coverslip facing the cortical surface was coated with poly-L-lysine (P4707; Sigma-Aldrich, Taufkirchen, Germany) and a sparse layer of 40 nm fluorescent beads (yellow-green FluoSpheres™, Thermo Fisher Scientific, Waltham, MA) to render it visible for fluorescence widefield, confocal and STED imaging. Finally, a thin layer of silicon polymer (First Contact; Photonic Cleaning Technologies, Platteville, WI) was applied to the outer side of the cover glass to protect the glass surface from dirt and scratches until the actual imaging commenced.

### I*n vivo* STED microscope and chronic *in vivo* imaging

We built a scanning STED microscope attached to an upright microscope stand (Leica Microsystems GmbH, Wetzlar, Germany), as previously described (*9, 47*). In brief, STED light was delivered by a Ti:Sapphire laser (MaiTai; Spectra-Physics, Santa Clara, CA), followed by an OPO (APE, Berlin, Germany), emitting 80MHz pulses at 595nm wavelength. The pulses were stretched to ~300ps by dispersion in a glass rod and a 120m long polarization-preserving fibre (OZ Optics, Ottawa, Canada). A helical phase delay of 0 to 2π was introduced by transmitting the STED beams through a vortex phase plate (RPC Photonics, Rochester, NY). For excitation a pulsed laser diode operating at 483nm, emitting pulses of 100ps duration (PiLas, Advanced Laser Diode Systems, Berlin, Germany) was used. After combining the excitation and STED beam via a dichroic mirror both beams were passing a Yanus scan head (Till Photonics-FEI, Gräfelfing, Germany), consisting of two galvanometric scanners and relay optics, and then were focused into the 1.3NA objective lens (PL APO, 63x, glycerol; Leica, Wetzlar, Germany). Temporal overlap was ensured by triggering the excitation laser diode with the STED light pulses. The back-projected fluorescent light was filtered with a 525/50nm band-pass and focused on a multimode fibre for confocal detection, connected to an avalanche photodiode detector (APD, Excelitas, Waltham, MA).

*In vivo* STED imaging was initiated upon a recovery period of 3-4 weeks after window implantation. Mice were anaesthetised as stated above. The head of the mouse was screwed to the tiltable head holder on the mounting plate. The window was aligned perpendicular to the optical axis of the microscope with the help of a home-built optical alignment device (*8*). The mouse was then moved to the microscope. Blood vessels on the cortical surface served as landmarks for the realignment of imaging spots. A confocal z stack, covering the dendrite of interest and the fluorescent beads adhering to the coverslip, allowed for the accurate determination of the depth of the image plane. Typically, imaging was performed at a cortical depth of 15-35 μm. The correction collar of the objective (PL APO, 63x, glycerol, 1.3NA; Leica, Wetzlar, Germany) was adjusted at each depth to compensate for spherical aberrations in the tissue in order to optimize the STED resolution. The excitation and STED laser power were kept at a minimum to avoid phototoxicity, typically evidenced as blebbing of neurites. After completion of the imaging session the anaesthesia was antagonized with Atipamezole (2.5mg/kg) and Buprenorphine (0.1mg/kg). *In vivo* STED imaging was conducted twice a week, i.e. at 3-4 day intervals.

Imaging parameters: GFP was excited with 4.5μW and depleted with an average STED power of 11.3-14mW at the back aperture of the objective. Z-stacks were recorded at 600nm increments and at a pixel size of 30nm x 30nm in x and y and a pixel dwell time of 5μs. Images with a signal >5Mc/s were corrected for the actual count rate according to the instructions of the manufacturer of the detector (APD, SPCM-AQRH-13).

### Image analysis

Spine morphology was analysed manually using Fiji (*48*). Only motion-artefact-free image stacks were selected for data analysis. Spines, emanating laterally of the parent dendrite and captured within one or two adjacent imaging frames and appearing in at least two consecutive time points were analysed. Dynamics of spine morphological parameters were assessed in stable spines. Spine neck length was measured by drawing a line along the neck from its base at the dendrite to the beginning of the spine head using the freehand line tool (Figure 3A). To compute the spine head size we measured the longest axis (a) of the spine head and perpendicular to that the shortest axis (b) of the spine head using the straight line tool. The size of the spine head cross section was calculated by estimating the area (A) by an ellipse: A=π(a/2*b/2). The spine neck width was measured as the full-width at half-maximum (FWHM) of a line profile (average of 3 lines) of the spine neck at its thinnest position.

Normalized relative changes at time point ‘t_x+1_’ were calculated to the previous time point ‘t_x_’ by computing (M(t_x+1_)-M(t_x_))/(M(t_x+1_)+M(t_x_)). ‘M’ stands for the measured spine parameter spine head size, neck length or neck width.

The covariance function (Cov) was computed for different time intervals Δt by

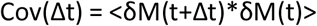

and the correlation function (Corr) by

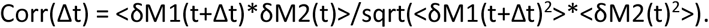

δ denotes the fluctuation of the mean: δM(t)= M(t)-<M(t)> and the brackets <…> the average over time and over the empirical distribution of spines. The auto-correlation was computed with M1 = M2 and the cross-correlation was computed with M1 and M2 denoting different spine parameters. The error bars refer to the bootstrapped standard deviation based on resampling the data 100 times.

The angle at which a spine emanated from the parent dendrite (neck dendrite angle) was measured using the angle tool in Fiji (note, this angle is limited to 90° as always the smaller angle is reported, Figure 3H). The angle formed between the spine neck and the spine head (neck head angle) was also measured using the angle tool (Figure 3H). Also here the smaller angle is reported, i.e. the maximum value of this angle 180°.

In order to assess overall spine density, all spines along the captured dendrites were counted over three consecutive time points.

### Statistics

Statistical analyses were performed either in MATLAB or GraphPad Prism. The statistical test and precision measure is specified in the figure legend together with the p-value. Significance was defined by *p<0.05; **p<0.01; ***p<0.001. The total number of mice, analyzed spines, number of dendrites per graph is summarized in supplementary table S1.

## General

We would like to thank Sandra Rode for help with Figure 1A.

## Author contributions

Conceptualization (SL, FW, KIW); Surgery (ACM, HS, VK, PS); STED microscope (KIW); Imaging (ACM, WW, KIW); Formal Analysis (Si.L, FW, KIW); Writing (SL, FW, KIW); Funding Acquisition (SL, KIW); Supervision (KIW).

## Data and materials availability

Requests for image data sets should be directed to corresponding authors and will be made available upon reasonable request. No material was generated.

