## Supplemental material for "Stable but not rigid: Long-term *in vivo* STED nanoscopy uncovers extensive remodeling of stable spines and indicates multiple drivers of structural plasticity"

### Supplementary Materials

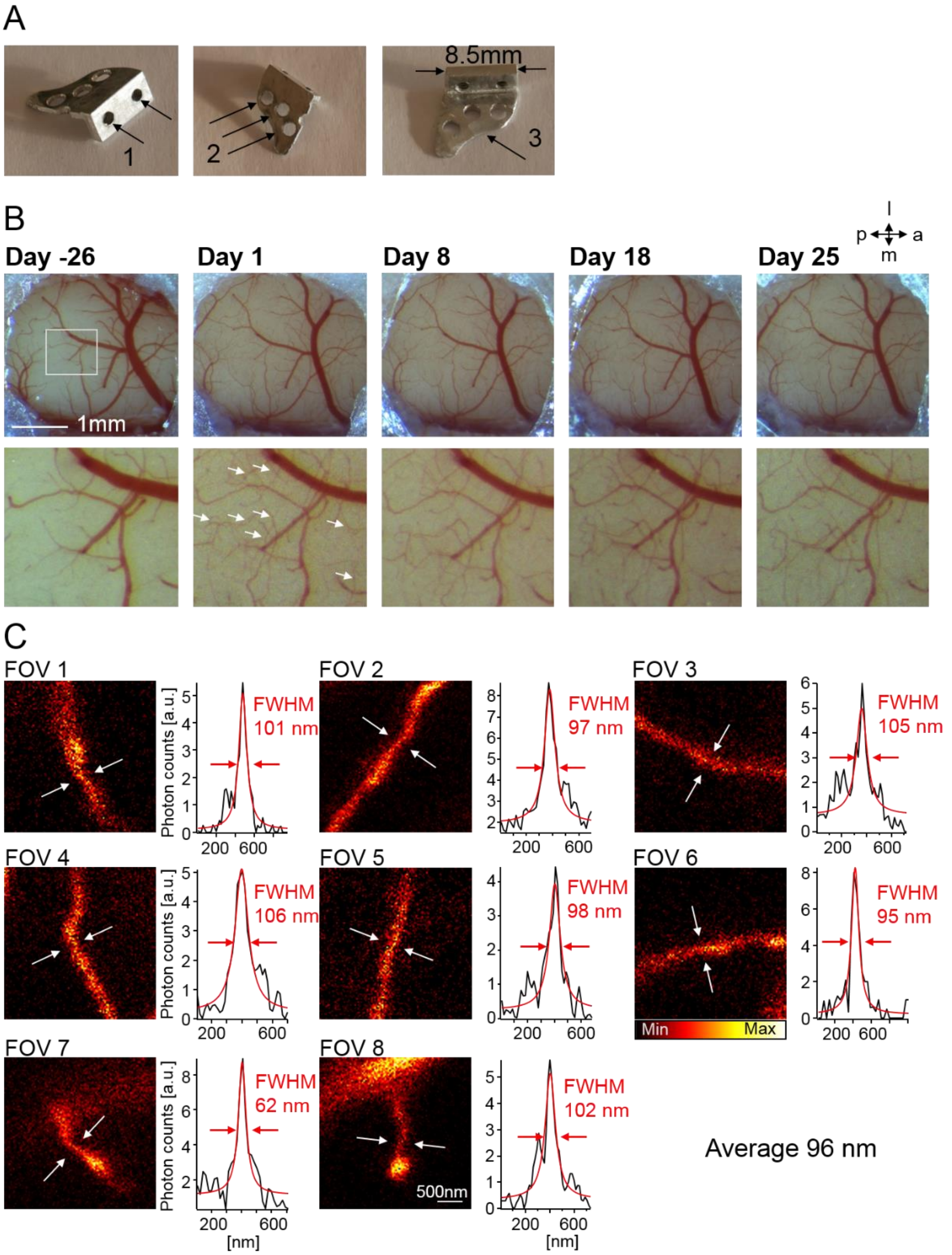


**Figure S1:** Cranial window for chronic STED imaging. (A) Head bar made of aluminium; threads for mounting (1), holes to apply resin (2) and curvature to fit skull (3). (B) Top: Motor cortex through implanted window at day of window implantation (day -26), start of the *in vivo* STED microscopy (day 1) and selected days of measurements. Bottom: Magnification of the marked area in top row. Dense network of sprouting new blood vessels at day 1 of imaging are marked with white arrows. (C) Estimation of STED microscopy resolution in chronic STED images. Magnified image details of raw STED microscopy images displaying axons or spines. Single frame. The line profile corresponds to an average of 3 lines, fitted with a Lorentzian function. The average resolution based on these full-width-half maximum (FWHM) measurements is 96 nm, which reflects an upper estimate of the resolution due to the relatively large size of axons. Abbreviations: FOV: Field of view; a: anterior, p:posterior, l: lateral, m: medial.

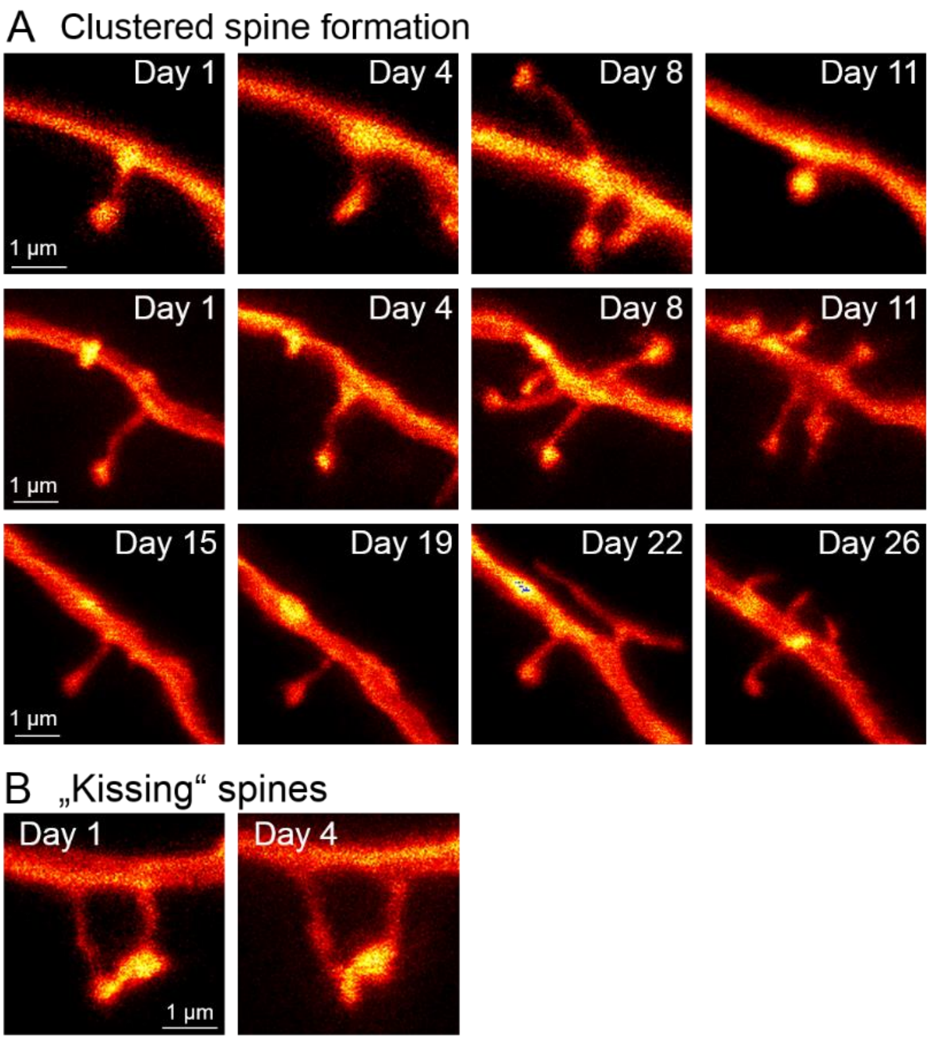


**Figure S2:** Selection of conspicuous spine shapes and dynamics. (A) Spines appear and disappear even in pairs (first and second line); spines grow and shrink over several µm within 3 days (third line). (B) Occasionally spine heads are in very close proximity.

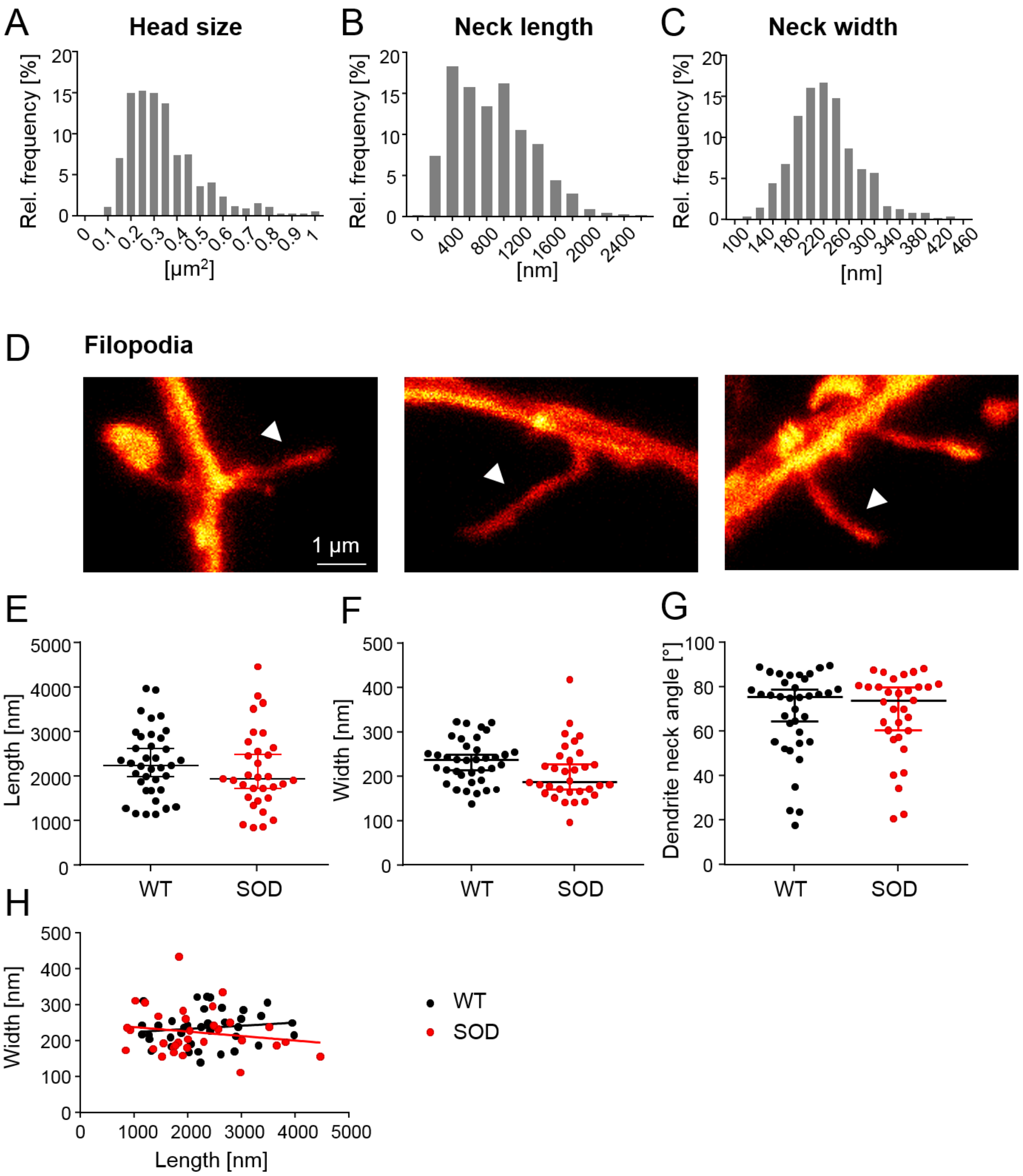


**Figure S3:** Distribution of (A) spine head sizes, (B) spine neck lengths and (C) neck cross sections in WT mice. (D-H) Filopodia analysis. (D) Representative images of filopodia (marked with arrow head). (E) Filopodia length ($p = 0.25$, KS test), (F) width ($p = 0.12$, KS test) and (G) dendrite to neck angle ($p = 0.99$, KS test) did not differ between WT and SOD mice. (H) Lack of correlation between width and length of filopodia (WT $p = 0.42$; SOD $p = 0.34$). Data are median +/- 95% CI (E–G). SOD: superoxide dismutase, mice harbouring the $SOD1^{G93A}$ mutation are a transgenic model of amyotrophic lateral sclerosis; WT: wild type.

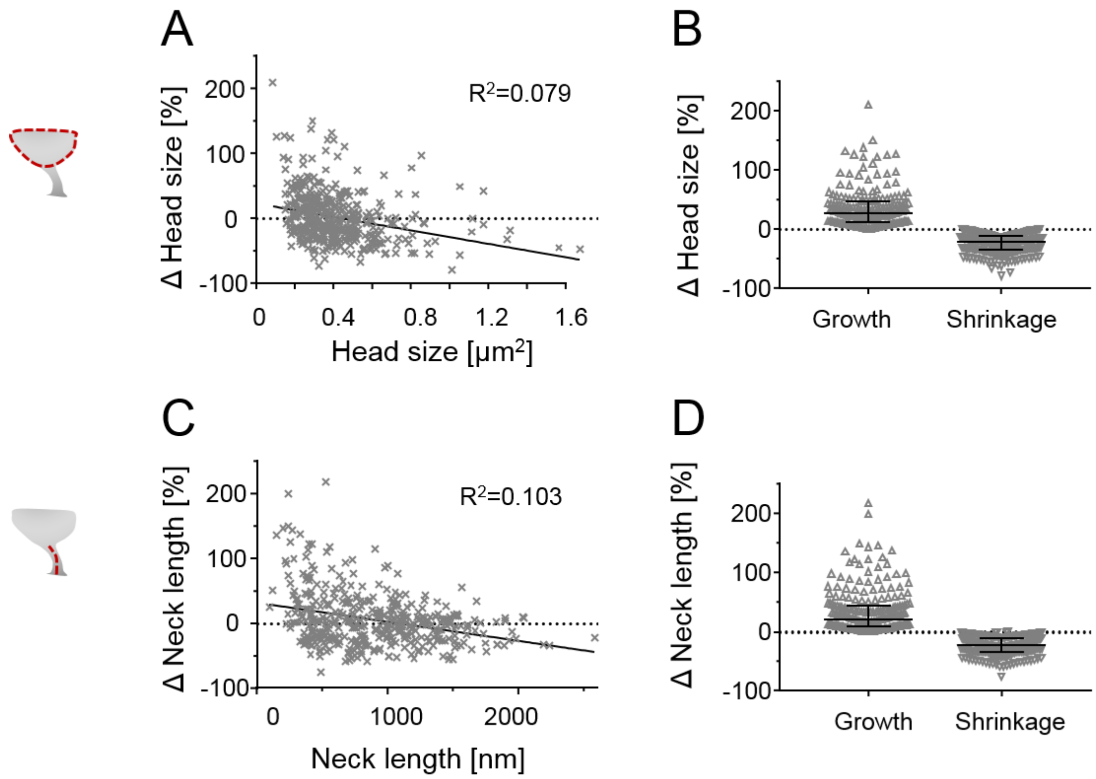


**Figure S4:** Changes of stable spine parameters in WT mice. (A) Percent change of head size as function of initial head size overlaid by linear regression ($p < 0.0001$). (B) Head size change for growing and shrinking spines in %. (C) Percent change of neck length as function of absolute neck length overlaid by linear regression ($p < 0.0001$). (D) Relative changes in neck length of extending and retracting spines in %. All changes assessed at 3-4 day intervals (A–D).

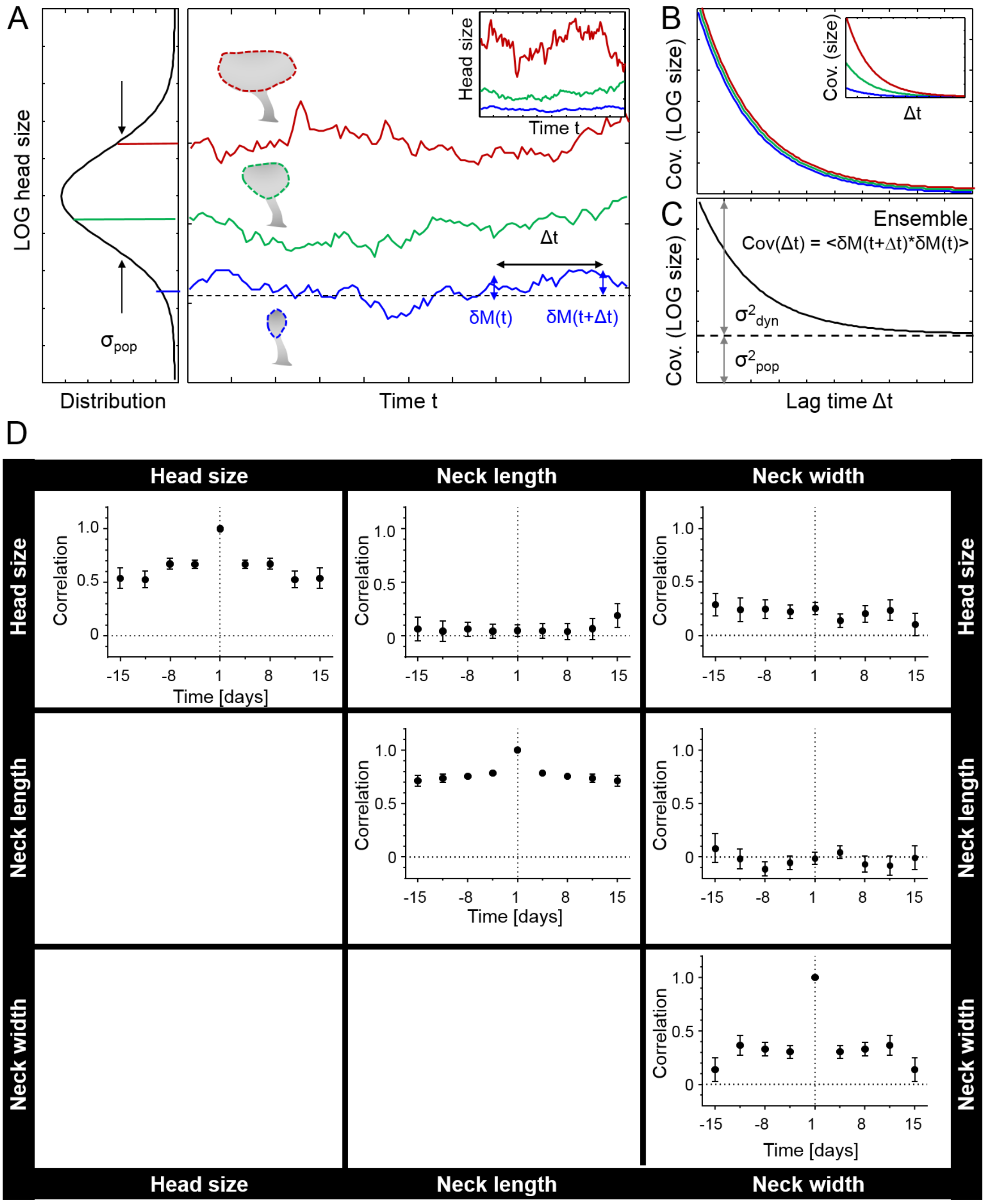


**Figure S5:** Covariance and correlation analysis of stable spines. (A-C) Covariance analysis of a model of dynamical fluctuations across a population of spine head size. (A) For multiplicative dynamics, the logarithm of the fluctuating quantity are approximately Gaussian distributed and exhibit uniform standard deviations $\sigma_{pop}$. Illustrated are the distribution of individual synapse mean values of head size (left) and model trajectories of fluctuations of the logarithm of the head size for three spines (small (blue), medium (green) and large (red)). Inset: On a linear scale, fluctuations of head size increase with increasing head size for multiplicative changes. (B) Temporal covariance functions for individual spines. δM deviation from mean value for different lag times Δt. (C) Ensemble covariance function as defined in Methods section. Double headed arrows indicate mean variance of dynamical fluctuations ($\sigma^2_{dyn}$) and variance of the population size distribution ($\sigma^2_{pop}$). (D) Auto- and cross-correlation of logarithmic values of measured spine parameters head size, neck length and neck width. Data are median and interquartile range (B, D) and SD of bootstrapped data (E).

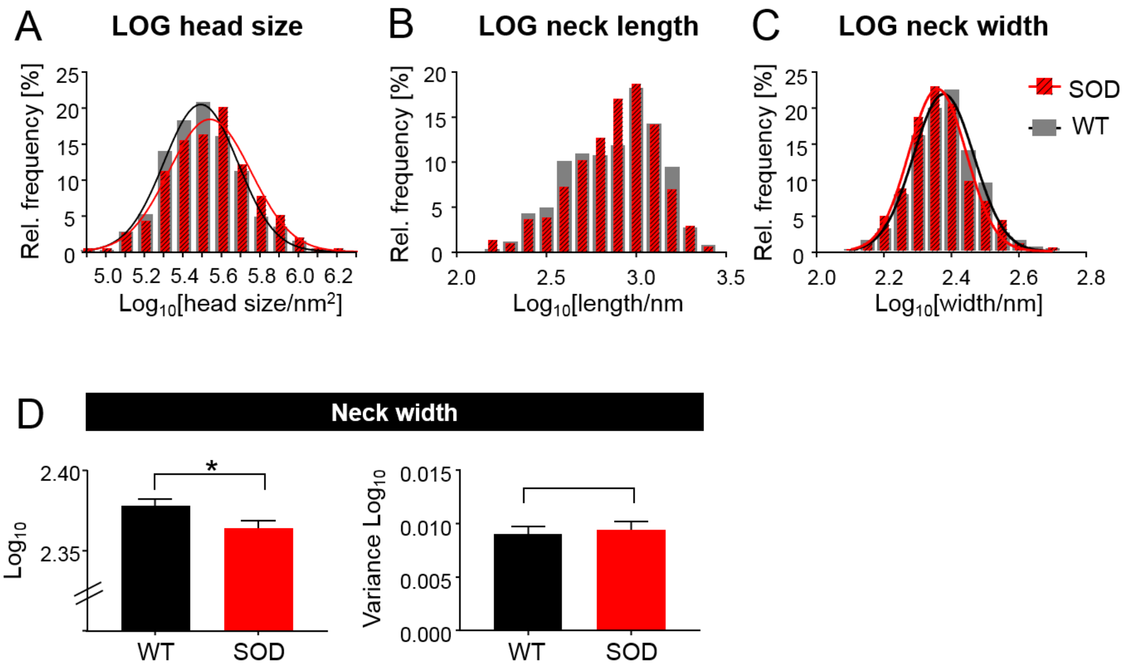


**Figure S6:** Distribution of $\log_{10}$ spine parameters. (A) Distribution of logarithmic spine head sizes overlaid by a gaussian fit for WT (grey) and SOD (red) mice. (B) Distribution of logarithmic neck length and (C) neck width. (D) Neck width of stable spines is lower in SOD compared to WT mice ($^{*}p = 0.037$, Unpaired t-test with Welch's correction), while changes in variance is not different ($p = 0.66$, F-test).

**Table S1:** Total number of analysed mice, field of views, dendrites and spines. NL: Neck length, HS: Head size, NW: neck width

| Figure | Number of mice | Number of field of views | Number of analysed values (spines or dendrites) |
|---|---|---|---|
| **3B** | 5 WT | 39 | 634 spines (pooled, all time points) |
| **3C** | 5 WT | 39 | 634 spines (pooled, all time points) |
| **3D** | 5 WT | 39 | 473 spines (pooled, all time points) |
| **3E** | 5 WT | 39 | 633 spines (pooled, all time points) |
| **3F** | 5 WT | 39 | 632 spines (pooled, all time points) |
| **3G** | 5 WT | 39 | 472 spines (pooled, all time points) |
| **3I** | 5 WT | 15 | 468 spines (pooled, 3 time points) |
| **3J** | 5 WT | 15 | 468 spines (pooled, 3 time points) |
| **3K** | 5 WT | 15 | 462 spines (pooled, 3 time points) |
| **4B** | 5 WT | 39 | |
| **4B, day 1** | | | NL=185, HS=185, NW=147 spines |
| **4B, day 4.5** | | | NL=160, HS=160, NW=108 spines |
| **4B, day 8** | | | NL=127, HS=127, NW=90 spines |
| **4B, day 11.5** | | | NL=64, HS=64, NW=48 spines |
| **4B, day 15** | | | NL=35, HS=35, NW=32 spines |
| **4B, day 18.5** | | | NL=36, HS=36, NW=29 spines |
| **4B, day 22** | | | NL=26, HS=26, NW=20 spines |
| **4B, day 24.5** | | | NL=10, HS=10, NW=9 spines |
| **4C-F** | 5 WT | 39 | 430 spines (pooled, all time points) |
| **4G-I** | 5 WT | 39 | |
| **4G-I, day 1** | | | NL=634, HS=634, NW=473 spines |
| **4G-I, day 4.5** | | | NL=423, HS=423, NW=257 spines |
| **4G-I, day 8** | | | NL=272, HS=272, NW=180 spines |
| **4G-I, day 11.5** | | | NL=154, HS=154, NW=100 spines |
| **4G-I, day 15** | | | NL=85, HS=85, NW=65 spines |
| **4K** | | 15 WT, 15 SOD | 111 stable, 71 lost, 66 gained spines |
| **4L** | | 15 WT, 15 SOD | 111 stable, 71 lost, 66 gained spines |
| **4M** | | 15 WT, 15 SOD | 111 stable, 71 lost, 66 gained spines |
| **5B** | | 15 WT, 15 SOD | 15 dendrites WT, 15 dendrites SOD |
| **5C** | 5 WT, 6 SOD | 39 WT, 38 SOD | 634 WT, 552 SOD (pooled) |
| **5D** | 5 WT, 6 SOD | 39 WT, 38 SOD | 634 WT, 552 SOD spines (pooled, all time points) |
| **5E Head size** | 5 WT, 6 SOD | 39 WT, 38 SOD | 430 WT, 394 SOD spines (pooled, all time points) |

| | | | |
|---|---|---|---|
| **5E Neck length** | 5 WT, 6 SOD | 39 WT, 38 SOD | 430 WT, 394 SOD spines (pooled, all time points) |
